# Population structure and genetic diversity of *Salmonella* Typhimurium in avian hosts

**DOI:** 10.1101/2022.11.17.516949

**Authors:** Yezhi Fu, Nkuchia M. M’ikanatha, Edward G. Dudley

## Abstract

Within-host evolution of bacterial pathogens can lead to host-associated variants of the same species or serovar. Identification and characterization of closely related variants from diverse host species are crucial to public health and host-pathogen adaptation research. However, the work remained largely underexplored at a strain level until the advent of whole-genome sequencing (WGS). Here, we performed WGS-based subtyping and analyses of *Salmonella enterica* serovar Typhimurium (*n* = 787) from different wild birds across 18 countries over a 75-year period. We revealed seven avian host-associated *S.* Typhimurium variants/lineages. These lineages emerged globally over short timescales and presented genetic features distinct from *S.* Typhimurium lineages circulating among humans and domestic animals. We further showed that, in terms of virulence, host adaptation of these variants was driven by genome degradation. Our results provide a snapshot of the population structure and genetic diversity of *S.* Typhimurium within avian hosts. We also demonstrate the value of WGS-based subtyping and analyses in unravelling closely related variants at the strain level.

## Introduction

*Salmonella enterica* is a major zoonotic bacterial pathogen that causes morbidity and mortality in humans and animals worldwide^1,2^. More than 2,500 serovars have been identified within *Salmonella enterica* according to the distinct combination of O and H antigens^3^. These serovars are roughly grouped into two categories based on their host specificity, i.e., serovars with broad host range (generalists) and narrow host range (specialists)^4^. *Salmonella enterica* serovar Typhimurium (*S.* Typhimurium) and *S.* Enteritidis are examples of generalists that can colonize and cause diseases in a wide variety of host species such as humans, livestock, poultry, and wildlife. However, *S.* Typhi and *S.* Paratyphi A are restricted to humans and higher primates^5,6^, while *S.* Choleraesuis (pig adapted), *S.* Dublin (cattle adapted), *S.* Abortusovis (sheep adapted), and *S.* Gallinarum (avian adapted) are associated with specific livestock or poultry^7^.

Although *S.* Typhimurium is considered the prototypical generalist serovar, epidemiologic evidence supports that this serovar has undergone adaptive evolution within specific host species, particularly in wild birds. Some avian host-associated *S.* Typhimurium variants identified by phage typing include definite phage type (DT) 2 and DT99 circulating in pigeon^8^, DT8 linked to duck/goose^9^, and DT40 and DT56 adapted to passerine^10^. Recently, we also documented three *S.* Typhimurium variants associated with larid, water bird, and passerine^11^. The emergence of host-associated variants in a broad-host-range serovar suggests that defining generalist bacterial pathogens at a species or serovar level is an oversimplification. It also highlights the importance of within-host evolution in shaping bacterial genetic diversity and host specificity.

Each host species represents a distinct ecological niche for bacterial pathogens. Over the course of colonization and infection, bacterial pathogens face challenges from the host species such as host immune response, antibiotic treatment, and native microbiota. Such challenges put selective pressure on bacterial pathogens and force them to evolve within the host^12^. As a result, bacterial pathogens are subjected to genomic changes to develop mechanisms of immune evasion and antimicrobial resistance (AMR), leading to emerging variants of the same species^13^. Wild birds constitute unique but underexplored ecological niches for microbes. Bacterial pathogens colonizing avian hosts may evolve divergently from their relatives residing in domestic animals due to difference in host environment (e.g., body temperature, immune system, exposure to antibiotics)^8,14,15^. Therefore, avian hosts may represent underestimated reservoirs for emerging pathogenic variants.

The emergence of new variants of bacterial pathogens poses a threat to public health as they may present distinct pathogenicity and epidemicity. It is important to identify new variants, characterize their genetic diversity, and correlate individual variants with their respective hosts. This will contribute to our understanding of the evolution, adaptability, and pathogenicity potential of bacterial pathogens within diverse hosts, and also be valuable for outbreak investigation and infection control/treatment. The traditional antibody-based serotyping method is used to differentiate between bacterial variants of the same species to a serovar level based on their surface antigens^3^. However, serotyping cannot distinguish bacterial variants of the same serovar. A variety of subtyping techniques such as pulsed-field gel electrophoresis (PFGE)^16,17^, seven-housekeeping-gene multilocus sequence typing (MLST)^18,19^, and phage typing^20^ have been developed for the latter purpose. Although these methods have been routinely used in surveillance for bacterial pathogens, they still lack resolution in discriminating between closely related variants at the strain level. Moreover, they cannot provide genetic information such as antimicrobial resistance and virulence of the tested variants^21^. The advance in whole-genome sequencing (WGS)-based subtyping and analyses provides superior resolution in identifying bacterial pathogens and unravelling their phylogenetic relationships and genetic makeup^22^. Currently, single nucleotide polymorphism (SNP) and whole genome or core genome-based MLST analyses are among the most commonly adopted WGS-based subtyping methods, which can differentiate bacterial pathogens at a strain level^23,24^.

In this study, we performed WGS-based subtyping and analyses of 787 *S.* Typhimurium isolates collected from diverse wild birds during 1946-2021 across 18 countries. The overall goal of this study is to reveal the population structure and genetic diversity of *S.* Typhimurium within avian hosts. By identifying distinct *S.* Typhimurium variants associated with avian hosts using WGS-based subtyping and analyses, our specific objectives are to: 1) gain insights into how within-host evolution of bacterial pathogens shapes their host specificity; 2) identify the evolutionary and genetic basis of *S.* Typhimurium adaptation to different host species; 3) assess the use of WGS-based subtyping and analyses in distinguishing between closely related variants (strain level) from multiple host species.

## Results

### Collection of *S.* Typhimurium isolates from avian hosts

A total of 787 *S.* Typhimurium isolates from avian hosts (avian hosts herein refers to wild birds, and do not include domestic poultry) were retrieved from EnteroBase on January 10, 2022 (Supplementary Data 1). The avian hosts were grouped into six categories based on bird type/phylogeny^25,26^ (Fig. 1 and Supplementary Data 1), i.e., passerine (order Passeriformes, also known as songbirds or perching birds; such as sparrow, finch, siskin, cardinal; *n* = 207), larid (order Charadriiformes; such as gull and tern; *n* = 138), duck/goose (order Anseriformes; *n* = 37), pigeon (order Columbiformes; *n* = 58), water bird (clade Aequornithes, such as cormorant, heron, pelican, stork; *n* = 154), and others (avian hosts without a designated bird type at EnteroBase or other bird types not mentioned above; *n* = 193). The collection contained historical and contemporary (1946-2021) isolates sampled from 18 countries across North America (*n* = 587), Europe (*n* = 124), Oceania (*n* = 52), Asia (*n* = 18), South America (*n* = 5), and Africa (*n* = 1) (Fig. 1). Of note, among the 787 genomes at EnteroBase, our group sequenced and uploaded 414 genomes (collection year: 1978-2019; collection location: 43 US states). Overall, our collection represented the most diverse collection of *S.* Typhimurium from avian hosts at EnteroBase as of the retrieval time.

**Fig. 1:**
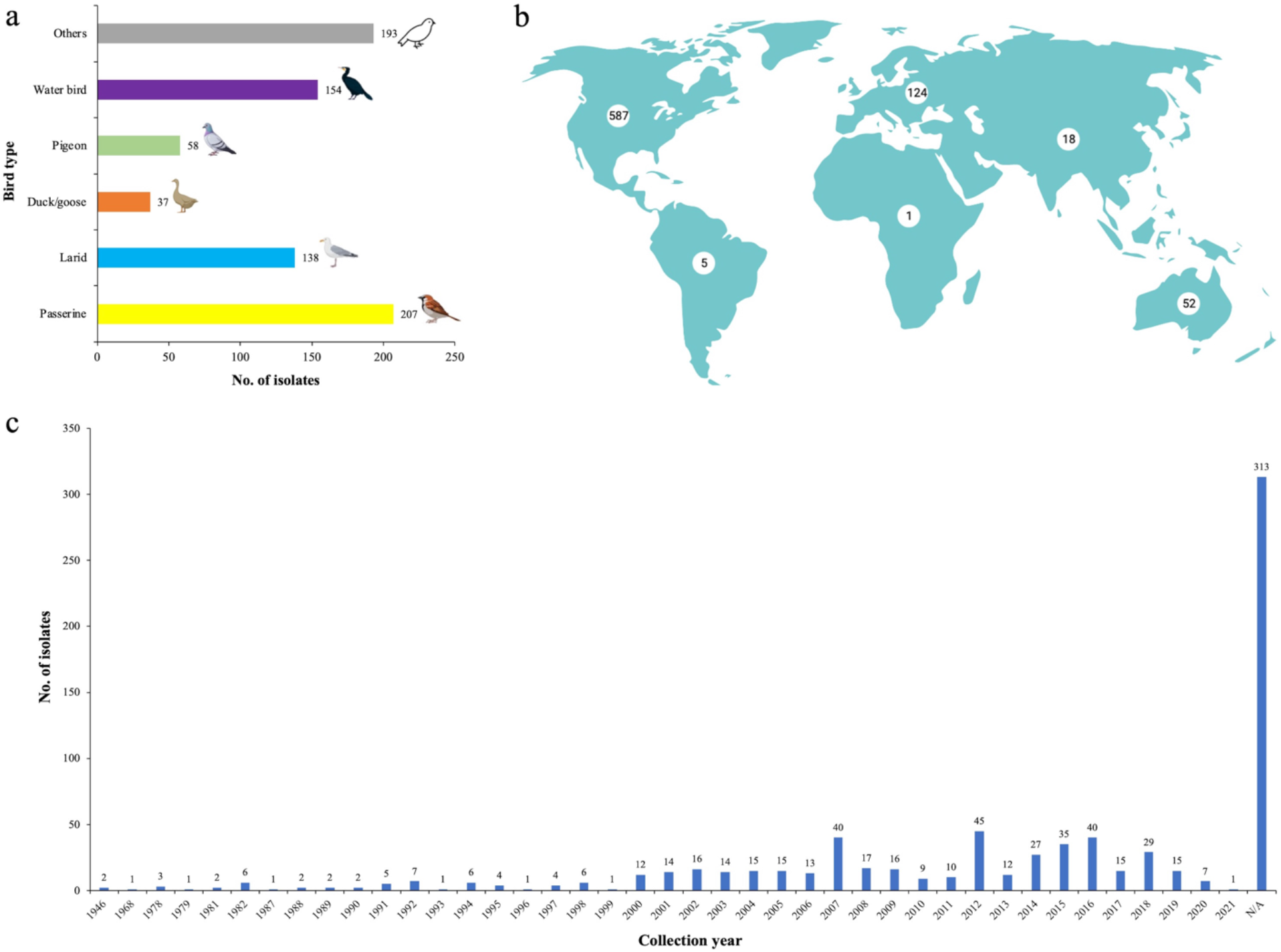
Avian isolates of *Salmonella* Typhimurium used in this study (*n* = 787). **a**, Number of isolates grouped by avian hosts. Bird type-others: avian hosts without a designated bird type at EnteroBase or any bird types not included in the defined categories. **b**, Number of isolates grouped by geographic locations. **c**, Number of isolates grouped by collection years. N/A: the collection year is not available.

### Population structure of *S.* Typhimurium from avian hosts

To investigate the population structure of *S.* Typhimurium from avian hosts, we generated a neighbor joining (NJ) tree of the 787 genomes (Fig. 2a) using the *Salmonella* wgMLST (whole genome MLST) scheme at EnteroBase. Ten *S.* Typhimurium lineages were present on the NJ tree, which included seven distinct lineages clustered by isolates (*n* = 633) primarily associated with specific bird types, i.e., passerine lineage 1 and lineage 2, larid lineage, duck/goose lineage, pigeon lineage 1 and lineage 2, and water bird lineage (Fig. 2a). The other three lineages on the NJ tree were formed by isolates from diverse bird types (Fig. 2a). As avian hosts usually are highly mobile and can migrate across different continents or countries, we also investigated the impact of geographic locations on the clustering pattern of the avian isolates. The seven *S.* Typhimurium lineages defined by bird type all contained isolates from ≥2 continents, indicating a global distribution of these lineages (Supplementary Fig. 1). Further, each individual lineage included isolates from multiple countries (Supplementary Fig. 2). Within the same lineage, isolates were observed to cluster based on collection countries. For examples, in passerine lineage 1, isolates from New Zealand clustered as a sublineage of the US passerine lineage (Supplementary Fig. 2a); in larid lineage, isolates from Australia clustered as a sublineage of the US larid lineage (Supplementary Fig. 2c). These observations indicate clonal expansions within different continents or countries, likely facilitated by bird migration^27,28^.

**Fig. 2:**
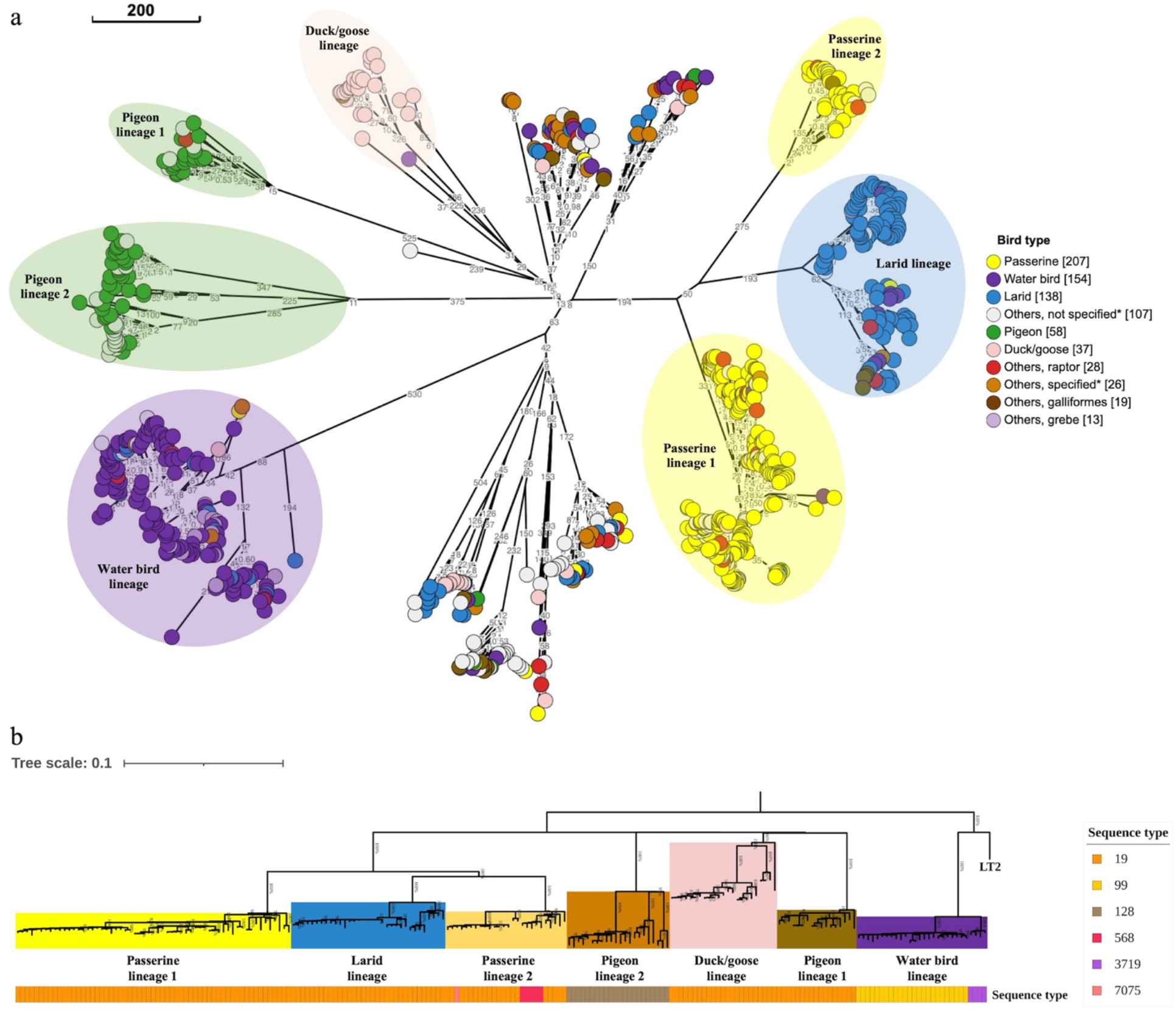
Population structure of globally sourced *Salmonella* Typhimurium isolates from avian hosts. **a**, Neighbor joining tree of the 787 *S.* Typhimurium isolates from avian hosts (https://enterobase.warwick.ac.uk/ms_tree?tree_id=70709). The NJ tree is constructed based on the *Salmonella* wgMLST scheme (21,065 loci) at EnteroBase. The scale bar indicates 200 wgMLST alleles. Allele differences between isolates are indicated by numbers on the connecting lines. In the legend “Bird type”, the number in brackets indicates the number of isolates from that specific bird type. “Other, not specified” represents avian hosts without a designated bird type at EnteroBase. “Other, specified” represents avian hosts that do not belong to passerine, larid, water bird, duck/goose, pigeon, and the number of isolates from these avian hosts is <10. More detailed information on individual bird type and its corresponding isolates can be found in Supplementary Data 1. **b**, Maximum-likelihood phylogenetic tree of the 207 *S.* Typhimurium isolates from avian hosts (See “Methods-Dataset collection” for the selection criteria for the 207 isolates out of the whole collection of 787 isolates). The tree is built based on 6,310 SNPs in the core genomic regions with reference to *S.* Typhimurium LT2 and rooted at midpoint. Individual avian host-associated lineages are supported by bootstrap value of 100%. The color strip “Sequence type” represents the *S.* Typhimurium multilocus sequence type determined by 7-gene (*aroC, dnaN, hemD*, *hisD*, *purE*, *sucA* and *thrA*) MLST.

We filtered the 787 genomes by excluding those without a collection year, location, specific bird host or other important metadata information. The filtered collection of 207 *S.* Typhimurium genomes (Supplementary Data 2) were used for further phylogenetic analysis and Bayesian inference. A maximum-likelihood (ML) phylogenetic tree based on 6,310 core-genome SNPs (cgSNPs) of the 207 genomes were built to validate the population structure of avian *S.* Typhimurium inferred by wgMLST. The lineages present in the cgSNP-based ML phylogenetic tree (Fig. 2b) were supported by robust bootstrap values of 100% and congruent with those formed in the NJ tree based on wgMLST.

A total of six STs (ST19, 99, 128, 568, 3719, and 7075) were identified among the seven lineages based on the classic seven-housekeeping-gene MLST method (Fig. 2b). Specifically, isolates from passerine lineage 1, larid lineage, duck/goose lineage, and pigeon lineage 1 all belonged to ST19, which is consistent with the fact that ST19 is one of the most prevalent *S.* Typhimurium sequence types detected in a broad range of hosts^19^. In addition, isolates from pigeon lineage 2 were represented by ST128, and variable STs were presented in isolates from passerine lineage 2 (i.e., ST19, 568, and 7075) and water bird lineage (i.e., ST99 and 3719). Therefore, sequence types defined by seven-housekeeping-gene MLST method did not distinguish between the lineages defined by bird type.

### Emergence times of avian *S.* Typhimurium lineages

Temporal signal of the sequence data was examined by TempEst^29^ before Bayesian molecular clock analysis. Moderate to strong temporal signals (correlation coefficient between 0.65 and 0.96) were detected in the sequence data (Supplementary Fig. 3). After confirming temporal signal, we built a Bayesian time-scaled phylogenetic tree using BEAST2 v2.6.5 to infer the emergence times of the lineages (Fig. 3). Based on Bayesian inference, passerine lineage 1, passerine lineage 2, larid lineage, duck/goose lineage, and pigeon lineage 1 emerged in ca. 1950 [95% highest probability density (HPD): 1940– 1959], ca. 1969 (95% HPD: 1959–1977), ca. 1943 (95% HPD: 1925–1957), ca. 1826 (95% HPD: 1771–1885), and ca. 1959 (95% HPD: 1947–1969), respectively (Fig. 3). Isolates from the five lineages mostly belonged to ST19 except that some isolates from passerine lineage 2 presented variable STs (Fig. 2b), indicating that these lineages diverged from a most recent common ancestor (MRCA) belonging to ST19. Pigeon lineage 2 (ST128) and water bird lineage (ST99 and 3719) evolved independently and formed in ca. 1847 (95% HPD: 1798–1886) and ca. 1953 (95% HPD: 1935–1967), respectively (Fig. 3). Of note, duck/goose lineage and pigeon lineage 2 emerged in 19^th^ century (i.e., 1826 for duck/goose lineage and 1847 for pigeon lineage 2), whereas the other five lineages formed within 20^th^ century during 1940-1970. The results show that *S.* Typhimurium evolved on short timescales to form individual lineages within avian hosts. We then estimated the median substitution rate for each lineage according to Bayesian inference. Median substitution rates for individual lineages ranged from 1.3 × 10^-7^ to 6.4 × 10^-7^ substitutions/site/year, with the lowest substitution rate for duck/goose lineage and the highest substitution rate for water bird lineage (Supplementary Fig. 4). These estimates are higher than the long-term (over million years) substitution rates in *Salmonella* and *E. coli* (10^-10^ to 10^-9^ substitutions per site per year)^30^, but similar to the short-term (over months or years) substitution rates reported for two ST313 lineages adapted to humans in sub-Saharan Africa (1.9 × 10^-7^ and 3.9 × 10^-7^ substitutions per site per year)^31^.

**Fig. 3:**
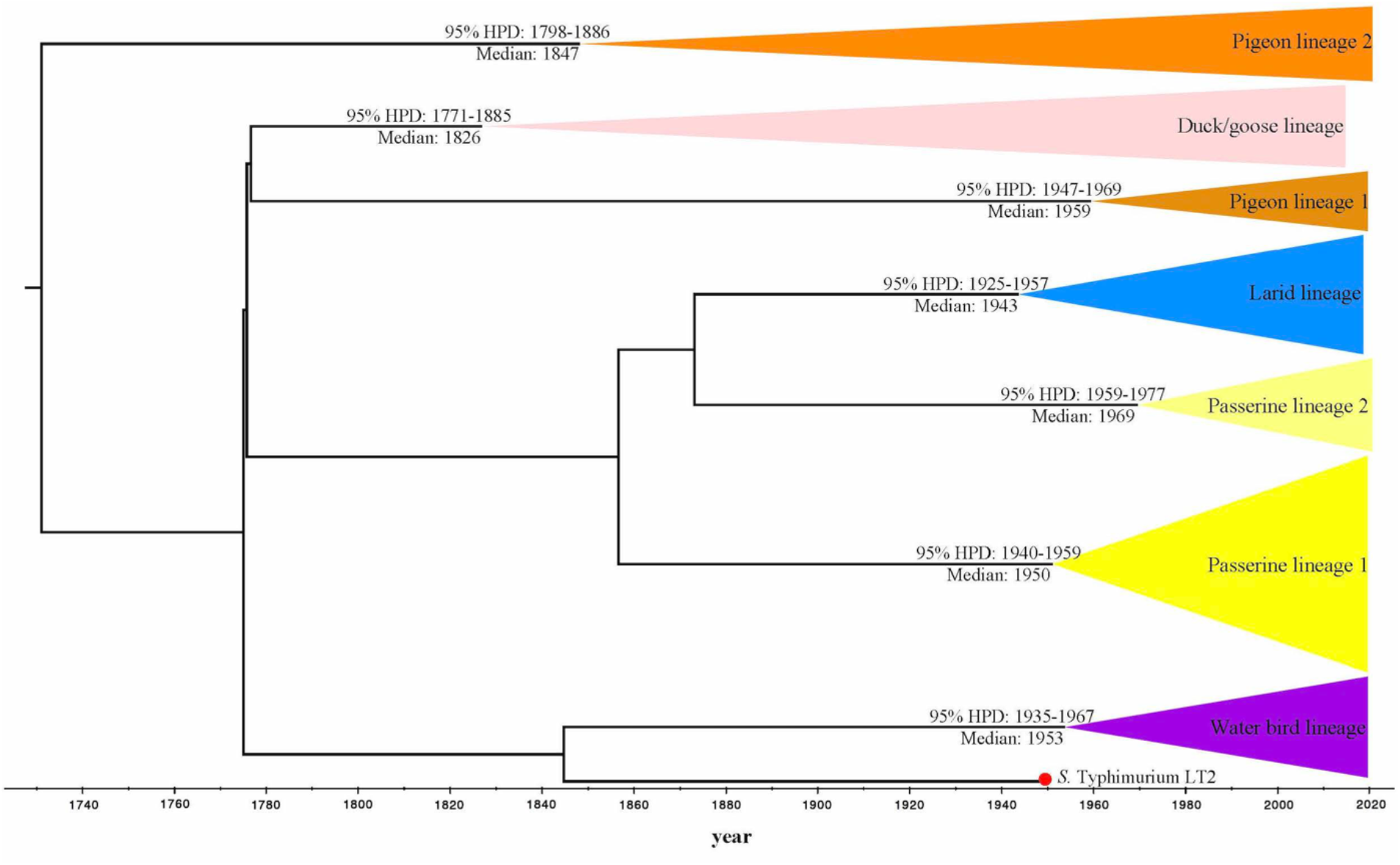
Emergence times of avian host-associated *Salmonella* Typhimurium lineages inferred by Bayesian time-scaled tree. Estimated emergence times of individual lineages are reported as median years with 95% highest posterior probability density (HPD). The red dot at the tree tip represents the reference genome from *S.* Typhimurium LT2 (collection year: ca. 1948). The posterior probability values of representative divergent events are >95% (not shown in the figure).

### Phylogenetic relationship of *S.* Typhimurium from avian and other diverse hosts

To investigate the phylogenetic relationship of avian isolates to other sourced isolates, we included 83 contextual genomes from diverse host species (humans, pigs, cattle, poultry) other than wild birds in the previous cgSNP-based ML phylogenetic tree. The contextual genomes represented the major epidemiologic *S.* Typhimurium lineages circulating globally (Supplementary Data 3). Taken together with the seven avian host-associated lineages, we presented a comprehensive population structure of *S.* Typhimurium in diverse hosts (Fig. 4). An NJ tree (Supplementary Fig. 5) of the 207 avian and 83 contextual genomes based on *Salmonella* wgMLST scheme at EnteroBase was built to complement the cgSNP-based ML phylogenetic tree. Isolates present in the NJ tree had the same clustering pattern with those shown in the ML phylogenetic tree based on cgSNPs (Fig. 4).

**Fig. 4:**
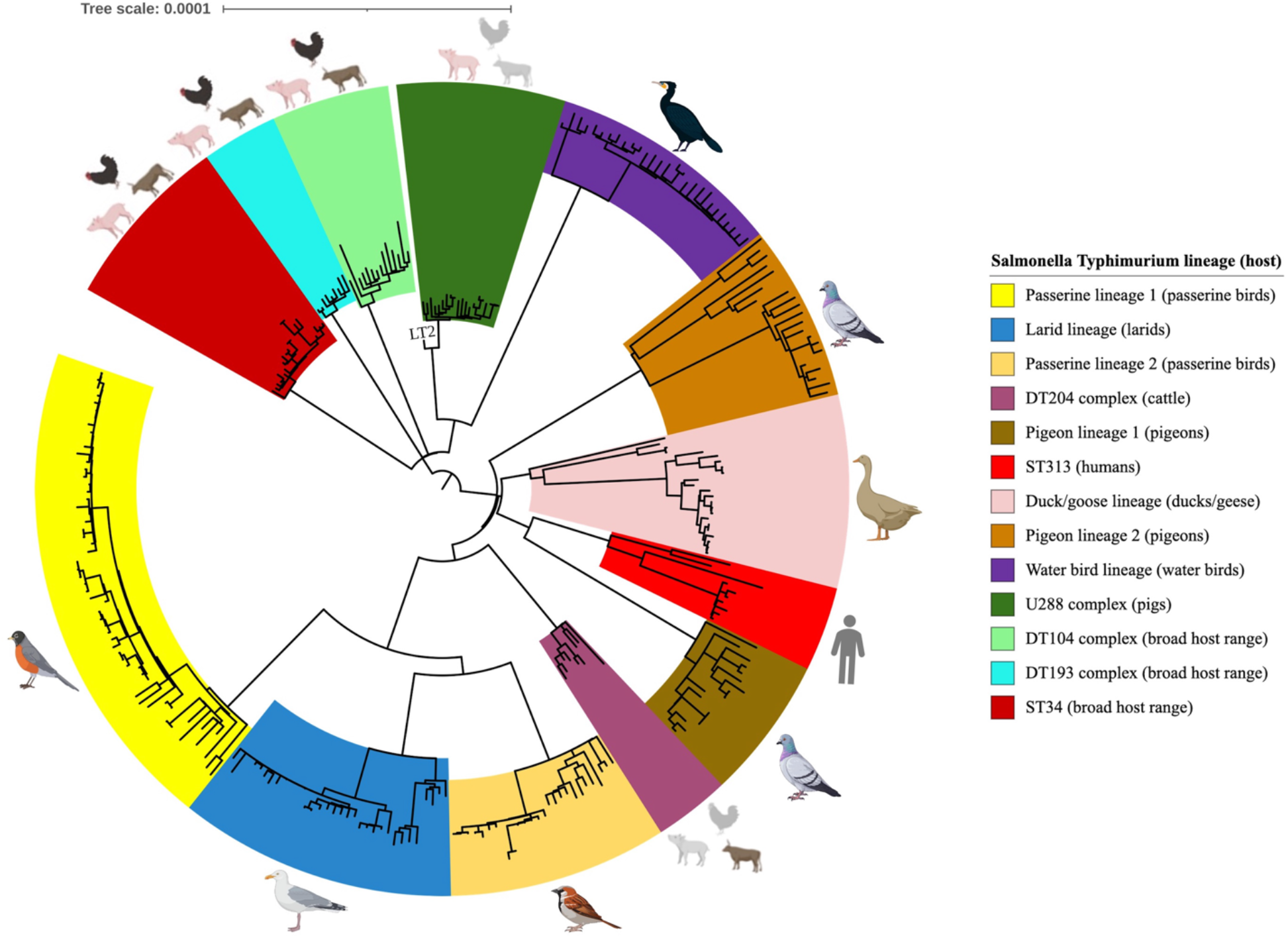
Phylogenetic relationship of *Salmonella* Typhimurium lineages circulating within diverse hosts (*n* = 290). The legend field at the right of the tree represents the *S.* Typhimurium lineage (primary host). Broad host range in parentheses indicates that isolates from the corresponding lineage are commonly identified among humans, cattle, pigs, and poultry. The specific host species in parentheses indicates that isolates from the corresponding lineage are primarily from that specific host. Individual lineages are correlated to their associated host species in the tree. Grey shaded host species in U288 complex lineage and DT204 complex lineage represent minor host other than primary host.

There were 13 lineages present in the ML phylogenetic tree (Fig. 4), which can be divided into two categories based on host range, i.e., lineages with broad host range (generalist lineages) and lineages with narrow host range (specialist lineages). Generalist lineages included monophasic *S.* Typhimurium ST34 lineage^32^, DT104 complex lineage^33^, and DT193 complex lineage^14^; on the other hand, specialist lineages contained DT204 complex lineage primarily associated with cattle^34^, U288 complex lineage possibly adapted to pigs^35^, human-adapted ST313 lineage causing invasive salmonellosis in sub-Saharan Africa^36,37^, and the seven lineages linked to specific bird types. By incorporating the host information into the cgSNP-based ML phylogenetic tree, we therefore were able to correlate individual lineages to specific host species (Fig. 4). It should be noted that generalist and specialist lineages are proposed in this study based on previous epidemiologic studies^9,14^, and lineages with narrow host range can occasionally infect other hosts^11,35,38–42^, indicating that adaptation of these lineages is still at an initial stage.

### Genomic comparison of *S.* Typhimurium lineages from avian and other diverse hosts

To explore the genetic diversity of *S.* Typhimurium variants, we performed comparative genomic analyses of the 207 genomes from avian hosts and 83 contextual genomes from other diverse hosts (humans, pigs, cattle, poultry). Pangenome analysis showed that the number of core genes (genes present in ≥99% isolates) shared by isolates within a specific lineage (henceforth referred to as lineage-associated core genes) ranged from 4,147 to 4,381, with the lowest being passerine lineage 1, and the highest being DT104 complex lineage (Fig. 5a; Supplementary Data 4). Isolates from all the 13 lineages shared 3,798 core genes, which we referred to as *S.* Typhimurium core genes. This number is smaller than previous estimates (3,836 or 3,910 core genes)^43,44^, possibly due to the increased genetic diversity in our dataset collection. By subtracting *S.* Typhimurium core genes from lineage-associated core genes, we calculated the number of core genes that represented a unique core-gene combination in a specific lineage (Fig. 5b). We further performed a pairwise comparison of lineage-associated core genes and found that individual lineages were differed from one another by an average number of 194 unique core genes (Supplementary Data 4). However, we did not find that avian host-associated lineages consistently presented much higher or lower number of unique core genes compared to lineages from other diverse hosts. In particular, passerine lineage 1 had the lowest average number of unique core genes (*n* = 123) relative to other lineages, while water bird lineage had the highest number (*n* = 265) (Supplementary Fig. 6).

**Fig. 5:**
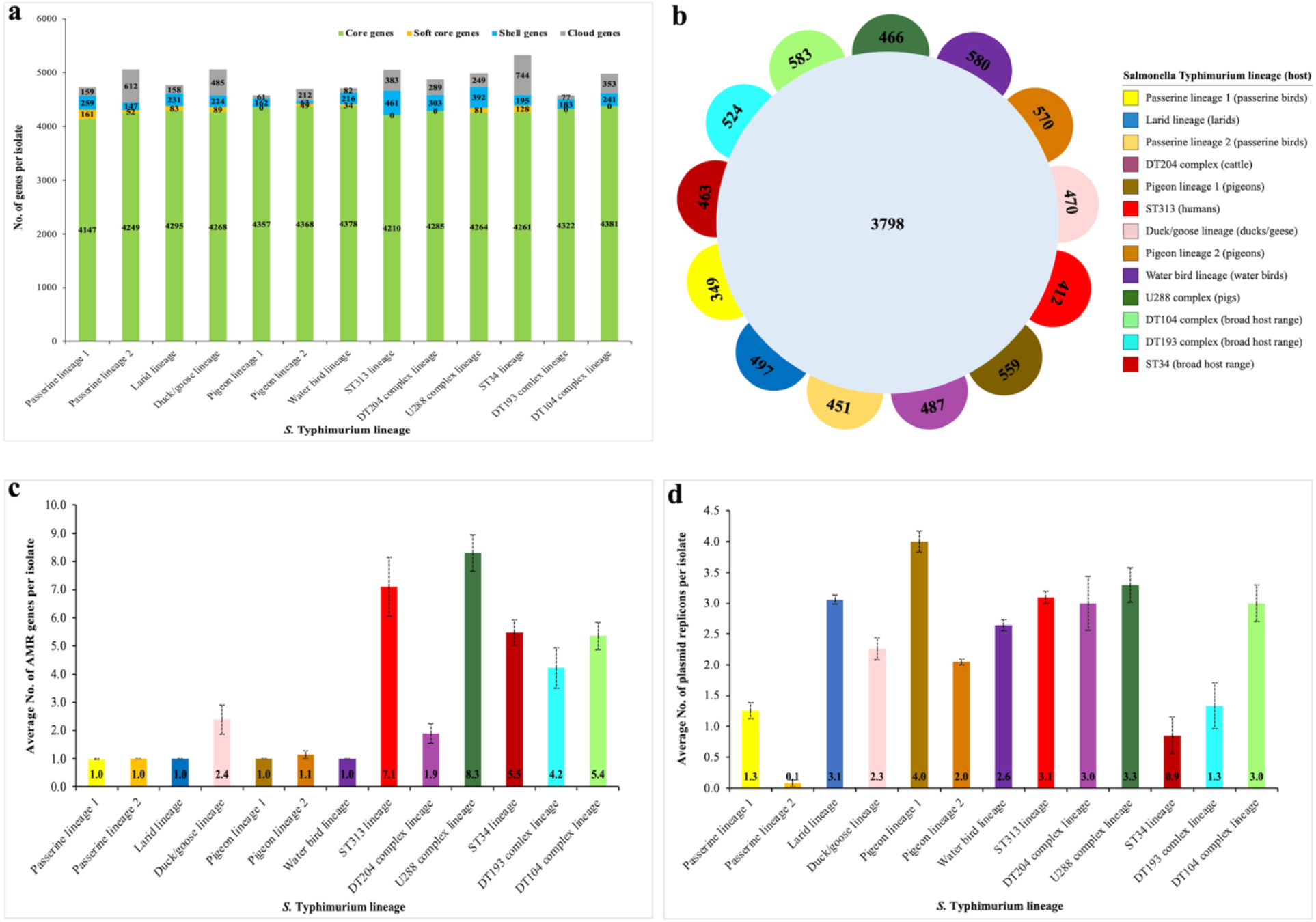
Genetic diversity of *Salmonella* Typhimurium lineages from diverse hosts (*n* = 290). **a**, Number of core genes (genes present in ≥99% isolates of the analyzed dataset), soft shell genes (genes present in 95%-99% isolates of the analyzed dataset), shell genes (genes present in 15%-95% isolates of the analyzed dataset), and cloud genes (genes present in 0%-15% isolates of the analyzed dataset) per isolate in individual lineages. **b**, Number of *S.* Typhimurium core genes (*n* = 3,798) and number of core genes that represent a unique core-gene combination in a specific lineage (see colored key). **c**, Average number of antimicrobial resistance (AMR) genes per isolate in individual lineages. **d**, Average number of plasmid replicons per isolate in individual lineages. The number of isolates in each lineage is: Passerine lineage 1 (*n* = 59); Passerine lineage 2 (*n* = 26); Larid lineage (*n* = 33); Duck/goose lineage (*n* = 23); Pigeon lineage 1 (*n* = 17); Pigeon lineage 2 (*n* = 21); Water bird lineage (*n* = 28); ST313 lineage (*n* = 10); DT204 complex lineage (*n* = 9); U288 complex lineage (*n* = 20); ST34 lineage (*n* = 21); DT193 complex lineage (*n* = 9); DT104 complex lineage (*n* = 14). Error bars represent standard error of the average number of a dataset.

AMR profiling revealed that all the isolates from avian host-associated lineages except duck/goose lineage lacked AMR genes (average number per isolate = 1) (Fig. 5c). The only AMR gene detected was *aac(6’)-Iaa* (Supplementary Data 5), which is a chromosomally encoded cryptic gene^45^. However, more AMR genes were detected in isolates from broad-host-range lineages (DT104, DT193, and ST34: average number per isolate >4), and lineages associated with humans (ST313: average number per isolate ≈ 7) or specific livestock (DT204 and U288: average number per isolate ≈ 2 and 8, respectively) (Fig. 5c).

Plasmid profiling revealed that most of the isolates from diverse lineages carried plasmid replicons IncFIB (70%; 203/290) and IncFII (74.5%; 216/290) (Supplementary Data 6) that belong to the *S.* Typhimurium-specific virulence plasmid pSLT^46^. However, both plasmid replicons were absent in all the isolates from passerine lineage 2 (*n* = 26) and ST34 lineage (*n* = 21) (Supplementary Data 6). As a result, isolates from the two lineages carried fewer plasmid replicons (average number per isolate <1) compared to isolates from other lineages (average number per isolate >1) (Fig. 5d). Additionally, isolates from passerine lineage 1 and DT193 complex lineage also tended to lose the two plasmid replicons (Supplementary Data 6). Specifically, both IncFIB and IncFII were absent in 40% (23/59) isolates from passerine lineage 1, while all the DT193 isolates (*n* = 9) lacked IncFIB and two DT193 isolates lacked IncFII (Supplementary Data 6).

### Prevalence of virulence-associated genome degradation in avian host-associated *S.* Typhimurium lineages

Our virulence profiling detected an average number of 114-116 virulence genes per isolate (Fig. 6a) for 9 out of the 13 lineages present on Fig. 4. The four lineages with fewer virulence genes per isolate were passerine lineage 1 (average number per isolate ≈ 113), passerine lineage 2 (average number per isolate ≈ 107), ST34 lineage (average number per isolate ≈ 107), and DT193 complex lineage (average number per isolate ≈ 113) (Fig. 6a). We further identified that the absent virulence genes were mostly encoded by pSLT, i.e., *pefABCD* (plasmid-encoded fimbriae), *rck* (resistance to complement killing), and *spvBCR (Salmonella* plasmid virulence) (Supplementary Data 7), which was consistent with the fact that isolates from these four lineages also completely or partially lacked pSLT replicons IncFIB or IncFII (Supplementary Data 6). For chromosomally encoded virulence genes, we only detected a complete loss of type 3 secretion system (T3SS) effector genes *gogB* in water bird lineage or *sopA* in DT193 complex lineage (Supplementary Data 7).

**Fig. 6:**
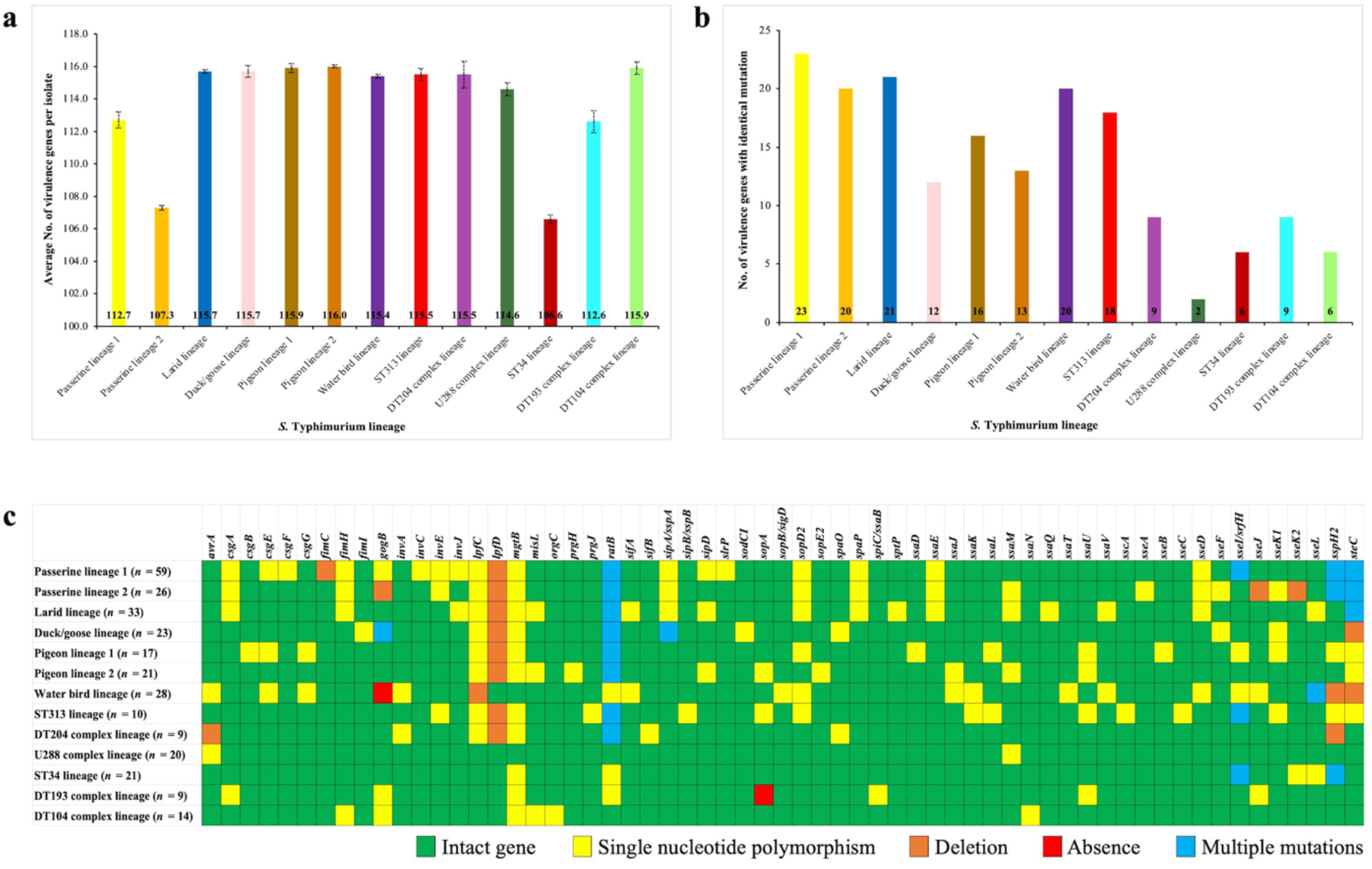
Virulence gene profiles of *Salmonella* Typhimurium lineages from diverse hosts (*n* = 290). **a,** Average number of virulence genes per isolate in individual lineages. Error bars represent standard error of the average number of a dataset. **b,** Number of virulence genes with identical mutation in individual lineages. **c**, Heatmap showing the mutation types of virulence genes in individual lineages. The number in parentheses indicates the number of isolates from that specific lineage. “Multiple mutations” indicates that several mutations occur in a virulence gene at different positions. The detailed mutation information (mutation type, mutation position, base-pair change) of each virulence gene in individual lineages can be found in Supplementary Data 8.

We also determined the number of chromosome-encoded virulence genes with identical mutation in individual lineages. Avian host-associated lineages and ST313 lineage adapted to humans had more than 10 (duck/goose lineage, pigeon lineage 1, pigeon lineage 2, ST313) or 20 (passerine lineage 1, passerine lineage 2, larid lineage, water bird lineage) identical mutant virulence genes; however, the number was less than 10 for lineages with broad host range (DT104, DT193, and ST34) or associated with livestock (DT204 and U288) (Fig. 6b). The types of mutation (Supplementary Data 8) were manually checked by aligning the virulence gene of interest against the reference virulence gene from *S.* Typhimurium LT2^47^ using BLAST. Among the 61 mutant chromosome-encoded virulence genes from different lineages, 47 were T3SS genes, five were curli genes (*csgA, csgB, csgE, csgF, csgG*), three were type 1 fimbriae genes (*fimC*, *fimH, fimI*), two were long polar fimbriae genes (*lpfC, lpfD*), and four were genes associated with other functions (*mgtB, misL, ratB, sodCl*) (Fig. 6c and Supplementary Data 8). The majority of mutations resulted from point mutations (SNPs) in T3SS genes, while a few virulence genes were subjected to deletions or multiple mutations (i.e., mutation occurs in more than one location in the gene) (Fig. 6c). We found that identical mutations in *lpfC* (substitution from C to T at position 328) and *lpfD* (deletion of GTTTGAGAAT at position 406-415) co-occurred in all specialist lineages except water bird lineage (single base-pair deletion in *lpfC* and intact *lpfD*) and U288 complex lineage (intact *lpfC* and *lpfD).* Each avian host-associated variants also had lineage-specific mutations. For instances, single base-pair deletion in *fimC* of passerine lineage 1, single base-pair deletions in *gogB, sseJ*, and *sseK2* of passerine lineage 2, SNPs in *sptP* and *ssaQ* of larid lineage, SNP in *sodCl* of duck/goose lineage, SNPs in *csgB, ssaD*, and *sseB* of pigeon lineage 1, SNPs in *prgH* and *sopE2* of pigeon lineage 2, and loss of *gogB* in water bird lineage (Fig. 6c).

## Discussion

Overall, our WGS-based subtyping and analyses identify seven avian host-associated *S.* Typhimurium lineages and provide new insights into the population structure and genetic diversity of *S.* Typhimurium from diverse host species (i.e., humans, livestock, poultry, wild birds). The avian host-associated lineages emerged over short timescales and present phylogenetic features (e.g., clustering based on bird type) and genetic traits (e.g., lack of AMR, lineage-specific virulence gene signatures) distinct from those formed by clinical isolates or isolates of domestic animal origin. Our findings suggest that some variants of this generalist serovar may be undergoing a convergent adaptive evolution driven by host species. From a virulence perspective, we find that genome degradation through point mutations (mainly SNPs) and deletions is the molecular basis of host adaptation of *S.* Typhimurium to avian hosts.

Among the 344,387 *Salmonella enterica* genomes deposited at EnteroBase as of January 10, 2022, only 0.5% of the genomes (*n* = 1,880) were from avian hosts. Our group sequenced and uploaded 699 out of the 1,880 genomes, which included 414 genomes of serovar Typhimurium. Therefore, our study makes a substantial contribution to the understanding of *S.* Typhimurium diversity with the identification of three new lineages associated with avian hosts (i.e., passerine lineage 1, larid lineage, and water bird lineage). Previous work on *S.* Typhimurium population structure focused on specific lineages formed by isolates from humans and domestic animals, and the geographic locations of these isolates were restricted to certain countries or regions^14,32,33,35–37,48^. As a result, the genetic diversity of this bacterial pathogen was underestimated due to a lack of representative isolates from wild animals, and the phylogenetic relationship of individual lineages remains largely unexplored on a global scale. Our study not only reveals the population structure of 787 avian isolates collected from 18 countries over a 75-year period, but explores the genetic diversity and phylogenetic relationship of globally sourced *S.* Typhimurium from diverse hosts.

As a prototype of generalist bacterial pathogens, *S.* Typhimurium can colonize and cause diseases in a variety of host species^49^. However, the identification of seven avian host-associated *S.* Typhimurium lineages indicates that some variants of serovar Typhimurium have adapted to specific avian host species. Previous studies also reported that DT204 and U288 complex lineages of *S.* Typhimurium were mainly restricted to cattle and pigs, respectively^34,35^, and ST313 lineage of *S.* Typhimurium was adapted to humans^36,37^. Therefore, it is more accurate to describe serovar Typhimurium as a collection of variants with different host range and degrees of host adaptation. The strong correlation of *S.* Typhimurium variants to specific hosts suggests that within-host evolution (host niche) is the primary driver in shaping host specificity of *S.* Typhimurium. Further, we did not find association of avian isolates with geographic locations at a lineage level. However, within some specific avian host-associated lineages, avian isolates from the same country clustered together to form a sublineage, indicating that geographic location (disperse limitation) serves as a less important evolutionary driver than host niche. The seven avian host-associated lineages emerged in 19^th^ and 20^th^ centuries (ca. 1826, 1847, 1943, 1950, 1953, 1959, and 1969, respectively), which occurred well after the divergence of avian host groups^25,26^. Similarly, the human-adapted ST313 sublineage L1, L2, and L3 in sub-Saharan Africa dated to around 1950, 1948, and 2007, respectively^37^. Collectively, these results support that host adaptation of *S.* Typhimurium is likely to be a relatively recent and ongoing process subjected to anthropogenic influence (e.g., globalization, antibiotic usage).

AMR profiles of *S.* Typhimurium lineages from diverse host species provide further evidence demonstrating the importance of host niches and anthropogenic activities in bacterial evolution. Our study shows that *S.* Typhimurium variants associated with avian hosts carried few AMR genes, while variants from humans or domestic animals had an average number of 2-8 AMR genes per isolate. Isolates evolve within avian hosts may be less likely to develop AMR as wild birds are rarely exposed to antibiotics in the natural environments; conversely, isolates from humans and domestic animals carry high number of AMR genes for the host species are frequently subjected to antibiotics, thus putting selective pressure on the colonized bacterial pathogens.

Genome degradation or loss-of-function mutation is a common pattern in adaptive evolution of *Salmonella*^50^. For example, loss or inactivation of fimbriae is linked to host adaptation^51,52^. Compared to host generalist serovar Enteritidis, host specialist serovars such as Dublin and Gallinarum accumulate more pseudogenes that lead to loss of fimbriae^50^. In this study, pseudogenization of the same fimbrial virulence gene network (*lpfC* and *lpfD*) due to deletion mutation was found in all specialist lineages except U288 complex lineage, suggesting inactivation of Lpf fimbriae may play an important role in transition of serovar Typhimurium from generalist to specialist. Additionally, it is reported that a group of T3SS effector proteins (SseL, SifB, SopD2, SseJ, SteB, SteC, SlrP, and SseK2) are frequently present in generalist serovars but lose functions in specialist serovars^53^. Similarly, we observed that more SNPs and deletions were accumulated in T3SS effector genes from host-associated lineages, which include but not limited to *sseL, sifB, sopD2, sseJ, steC, slrP*, and *sseK2.* It is likely that allelic variations in these T3SS effector genes may contribute to host specificity of *S.* Typhimurium.

A limitation of this study is the scarcity of *S.* Typhimurium isolates from avian hosts. Current WGS-based surveillance of bacterial pathogens primarily focuses on isolates from clinical samples, food samples, livestock, and poultry; however, isolates from wildlife have not been routinely collected and sequenced. As indicated in this study, wild animals such as wild birds represent remarkable but less studied reservoirs for emerging variants of bacterial pathogens. Epidemiologic studies have also revealed a correlation between some human and avian salmonellosis outbreaks, suggesting transmission of bacterial pathogens between wild birds and humans^38–42^. Although such transmission is rare relative to transmission between humans and humans or between humans and domestic animals^54,55^, they can still have a substantial impact on global health as avian hosts are highly mobile and possibly carry and spread bacterial pathogens over large distances^27,28^. In a One Health framework, current surveillance of bacterial pathogens needs to be not only focused on clinical isolates or isolates from domestic animals, but those originating from wild animals. We also note that the sequencing data in our collection is skewed toward *S.* Typhimurium isolates from North America, followed by Europe and Oceania, which is consistent with the fact that WGS has been widely used by countries (e.g., the United States, the United Kingdom, Australia) from these continents for surveillance of bacterial pathogens^56^. However, the state-of-the-art technology is less adopted in Asia, Africa, and South America, mostly due to economic reason^57^. Emerging epidemic lineages of bacterial pathogens may be circulating in these countries but underrepresented in current public repositories. Therefore, a global research collaboration is required to generate a robust and informative set of sequencing data to represent bacterial pathogens and their variants that cause diseases worldwide.

In conclusion, we reveal the population structure and genetic diversity of *S.* Typhimurium in avian and other diverse hosts. Our results indicate that within-host evolution has resulted in multiple host-associated *S.* Typhimurium lineages, which present genetic traits distinct from lineages with broader host range. Although our WGS-based subtyping and analyses are focused on serovar Typhimurium, the approach is translatable to other bacterial pathogens. It is expected that other generalist *Salmonella* serovars or bacterial pathogens such as *E. coli* and *Campylobacter* spp. commonly colonizing wild birds may have also undergone a similar adaptive evolution within avian hosts. Identifying these emerging host-associated variants and understanding the genetic basis of host adaptation will facilitate epidemiologic investigation, provide insight into the pathogenicity potential of the strain, and help design effective infection treatment/control strategies. For example, the lineage-specific mutations in virulence genes of avian host-associated lineages can serve as genetic markers for source tracking, and lack of AMR genes in avian host-associated *S.* Typhimurium variants means that antibiotics may treat the infection. Further, genome degradation in virulence genes may attenuate the pathogenicity of these variants to humans, making them of potential interest to study as vaccine candidates.

## Methods

### Dataset collection

*S.* Typhimurium genomes from avian hosts (*n* = 787) retrieved from EnteroBase (search term: source niche-wild animal; source type-avian; predicted serotype: serovar Typhimurium) were used to infer the population structure of this bacterial pathogen in wild birds. The avian isolates were collected over broad spatial and temporal scales (Supplementary Data 1). Among the 787 genomes deposited at EnteroBase, we sequenced and uploaded 414 genomes as part of a nationwide project collaborating with the US Geological Survey-National Wildlife Health Center to reveal antimicrobial resistance profile and evolutionary history of avian *S.* Typhimurium in the United States^11,58^. The *S.* Typhimurium isolates were collected from diseased or dead birds in 43 US states during 1978-2019 (Supplementary Data 1). The other 373 genomes were collected between 1946 and 2021 from 18 countries (including the United States) across the world and were publicly available at EnteroBase (Supplementary Data 1). We further refined the 787 genomes by excluding those without a designated collection year, location, bird host or those not belonging to an avian host-associated lineage. The filtered collection (*n* = 207) (Supplementary Data 2) was used for cgSNP-based ML phylogenetic analysis and Bayesian inference. In addition, contextual genomes (Supplementary Data 3; *n* = 83) from major *S.* Typhimurium epidemic lineages circulating worldwide were selected to infer the phylogenetic relationship and compare the genomic differences of *S.* Typhimurium from avian and other diverse host species (humans, livestock, poultry).

### DNA extraction and whole-genome sequencing

For DNA extraction of the avian isolates, each isolate was streaked onto xylose lysine deoxycholate agar plates and incubated for 18 h at 37 °C. A single colony was then picked, transferred to Luria-Bertani broth, and cultured overnight at 37 °C with continuous agitation (250 rpm). Genomic DNA was extracted using the Qiagen DNeasy® Blood & Tissue kit (Qiagen, Valencia, CA, USA) following the manufacturer’s instructions. DNA purity (A260/A280 ≥1.8) was confirmed using NanoDrop™ One (Thermo Scientific™, DE, USA) and DNA concentration was quantified using Qubit® 3.0 fluorometer (Thermo Fisher Scientific Inc., MA, USA). Extracted genomic DNA was stored at −20 °C before WGS. For WGS, DNA library was prepared using the Nextera XT DNA Library Prep Kit (Illumina Inc., San Diego, CA, USA), normalized using quantitation-based procedure, and pooled together at equal volume. The pooled library (600 μL) was denatured and sequenced on an Illumina MiSeq sequencer (Illumina Inc., San Diego, CA, USA).

### Quality assessment for raw reads

The quality of raw reads obtained in this study and downloaded from EnteroBase was assessed using the MicroRunQC workflow in GalaxyTrakr v2^59^. Sequence data passing quality control thresholds (i.e., average coverage >30, average quality score >30, number of contigs <400, total assembly length between 4.4 and 5.l Mb) were used for subsequent genomic analyses.

### Phylogenetic analysis

An NJ tree (https://enterobase.warwick.ac.uk/ms_tree?tree_id=70709) was built based on the wgMLST scheme (21,065 loci) at EnteroBase^60^ to infer population structure of *S.* Typhimurium from avian hosts (*n* = 787). Seven avian host-associated lineages were identified in the NJ tree. Genomes from the seven avian host-associated lineages were then refined as described in “**Data collection**”. The filtered collection of 207 *S.* Typhimurium genomes (Supplementary Data 2) was used to build the cgSNP-based ML phylogenetic tree. Specifically, Snippy (Galaxy v4.5.0) (https://github.com/tseemann/snippy) was used to generate a full alignment and find SNPs between the reference genome LT2 (RefSeq <u>NC_003197.1</u>) and the genomes of avian isolates, and Snippy-core (Galaxy v4.5.0) (https://github.com/tseemann/snippy) was used to convert the Snippy outputs into a core SNP alignment. The resultant core SNP alignment (6,310 SNPs in the core genomic regions) was used to construct a cgSNP-based ML phylogenetic tree by MEGA X v10.1.8 using the Tamura-Nei model and 1,000 bootstrap replicates^61^. Sequence types of the filtered *S.* Typhimurium isolates was identified using 7-gene (*aroC, dnaN, hemD, hisD, purE, sucA* and *thrA*) MLST at Enterobase^60^ and annotated on the cgSNP-based ML phylogenetic tree. We also added contextual genomes (Supplementary Data 3; *n* = 83) that represented the major *S.* Typhimurium epidemic lineages circulating globally in the cgSNP-based ML phylogenetic tree to infer the genetic relationship of lineages formed by avian and non-avian (e.g., humans, livestock, poultry) isolates. The cgSNP-based ML phylogenetic trees generated in this study were visualized and annotated using the Interactive Tree of Life (iTOL v6; https://itol.embl.de).

### Bayesian inference

The temporal signal of the sequence data was examined using TempEst v1.5.3^29^ before phylogenetic molecular clock analysis. Subsequently, a Bayesian time-scaled phylogenetic tree was constructed via BEAUti v2.6.5 and BEAST2 v2.6.5^62^ using the core SNP alignment (6,310 SNPs in the core genomic regions) generated from filtered collection (*n* = 207).

The parameters in BEAUti v2.6.5 were set as followings: Prior assumption-coalescent Bayesian skyline; Clock model-relaxed clock log normal; Markov chain Monte Carlo (MCMC): chain length-250 million, storing every 1,000 generations. Two independent runs with the same parameters were performed in BEAST2 v2.6.5 to ensure convergence. The resultant log files were viewed in Tracer v1.7.2 to ensure that the effective sample size (ESS) of key parameters was more than 200. A maximum clade credibility tree was created using TreeAnnotator v2.6.4 with burnin percentage of 10%. Finally, the tree was visualized using FigTree v1.4.4 (https://github.com/rambaut/figtree/releases) and annotated with the emergence times and substitution rates of individual lineages. To determine the substitution rate, we multiplied the substitution rate estimated by BEAST2 platform by the number of cgSNPs (6,310 bp), and then divided the product by the average genome size of the avian isolates (4,951,383 bp).

### Pangenome analysis

Raw reads of the 207 avian isolates and 83 contextual isolates from diverse host species were *de novo* assembled using Shovill (Galaxy v1.0.4)^63^ and then annotated by Prokka (Galaxy v1.14.6)^64^. The annotated contigs in GFF3 format produced by Prokka were taken by Roary (Galaxy v3.13.0)^65^ to calculate the pangenome with a minimum percentage identity of 95% for BLASTP. Specifically, lineage-associated core genes (i.e., genes present in more than 99% isolates from a specific lineage) were calculated by using genomes from individual lineages as input (passerine lineage 1: *n* = 59; passerine lineage 2: *n* = 26; larid lineage: *n* = 33; duck/goose lineage: *n* = 23; pigeon lineage 1: *n* = 17; pigeon lineage 2: *n* = 21; water bird lineage: *n* = 28; ST313 lineage: *n* = 10; DT204 complex lineage: *n* = 9; U288 complex lineage: *n* = 20; ST34 lineage: *n* = 21; DT193 complex lineage: *n* = 9; DT104 complex lineage: *n* = 14). *S.* Typhimurium core genes (i.e., genes present in more than 99% isolates from all lineages) were calculated by using genomes from all the isolates (*n* = 290) of avian and non-avian origin as input. We also performed a pairwise comparison of lineage-associated core genes to evaluate the genetic relatedness of individual lineages: First, the number of core genes shared by two lineages was calculated by using genomes from the two lineages; second, the number of core genes that differed the two lineages was obtained by subtracting the core genes shared by two lineages from lineage-associated core genes.

### AMR, plasmid, and virulence profiling

ABRicate (Galaxy v1.0.1)^66^ was used to identify the AMR genes, plasmid replicons, and virulence factors by aligning each draft genome assembly (see “**Pangenome analysis**”) against the ResFinder database^67^, PlasmidFinder database^68^, and Virulence Factor Database (VFDB)^69^, respectively. For all searches using ABRicate, minimum nucleotide identity and coverage thresholds of 80% and 80% were used, respectively. Virulence genes that were not 100% identical or covered with the reference virulence gene from VFDB may have deletions, insertions, or substitutions of interest. We then manually checked the mutation type by aligning the virulence gene of interest against the reference virulence gene from VFDB using BLAST (https://blast.ncbi.nlm.nih.gov/Blast.cgi).

## Data availability

Sequence data of the *S.* Typhimurium isolates from our lab (isolate name in the format “PSU-4 digits”, e.g., PSU-2718) are deposited in the NCBI Sequence Read Archive (SRA) (https://www.ncbi.nlm.nih.gov/sra) under BioProject <u>PRJNA357723</u>. Publicly available sequence data were downloaded from EnteroBase (https://enterobase.warwick.ac.uk/), NCBI SRA (https://www.ncbi.nlm.nih.gov/sra), and the European Nucleotide Archive (https://www.ebi.ac.uk/ena). Accession numbers of the genomes used in this study are listed in Supplementary Data 1-3.

## Acknowledgements

We thank David Hewitt for editing this manuscript prior to publication. This work is supported by the US Food and Drug Administration (Grant No. 1U19FD007114-01), US Department of Agriculture (Grant No. PEN4522), and Penn State College of Agricultural Sciences.

## Contributions

Y.F. designed the study, sequenced the US wild bird isolates, collected the globally sourced sequence data from EnteroBase, NCBI, and EMBL-EBI, performed the bioinformatics analyses of the data, interpreted the data, and wrote the draft manuscript; N.M.M. and E.G.D. contributed to interpretation of the data and manuscript revision.

## Corresponding author

Correspondence to Yezhi Fu and Edward G. Dudley.

## Ethics declarations

The authors declare no competing interests.

---

**Supplementary Fig. 1:**
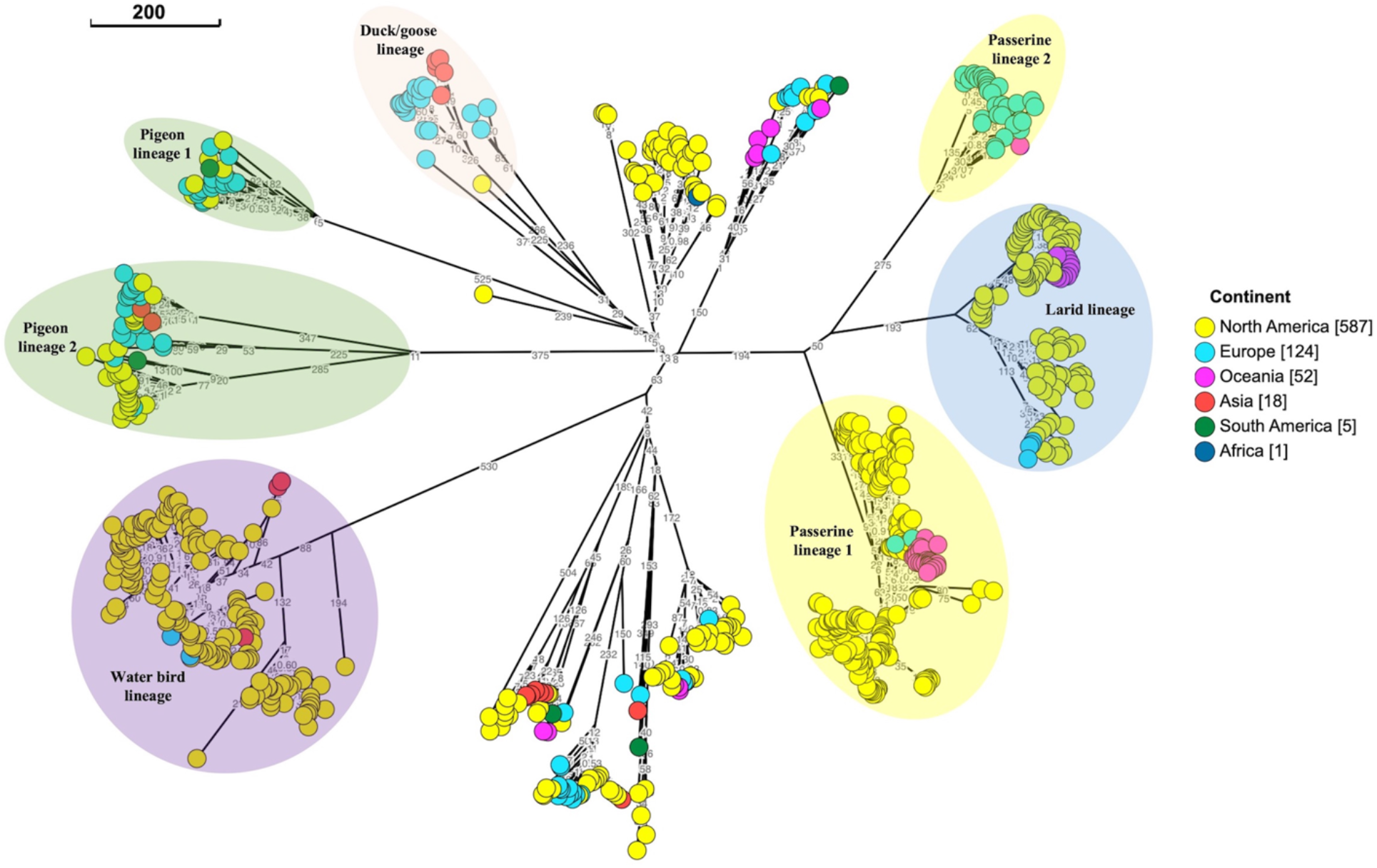
Neighbor joining tree based on wgMLST showing the geographic distribution of avian *Salmonella* Typhimurium isolates (*n* = 787) in six continents. In the legend “Continent”, the number in brackets indicates the number of isolates from that specific continent. The scale bar indicates 200 wgMLST alleles. Allele differences between isolates are indicated by numbers on the connecting lines.

**Supplementary Fig. 2:**
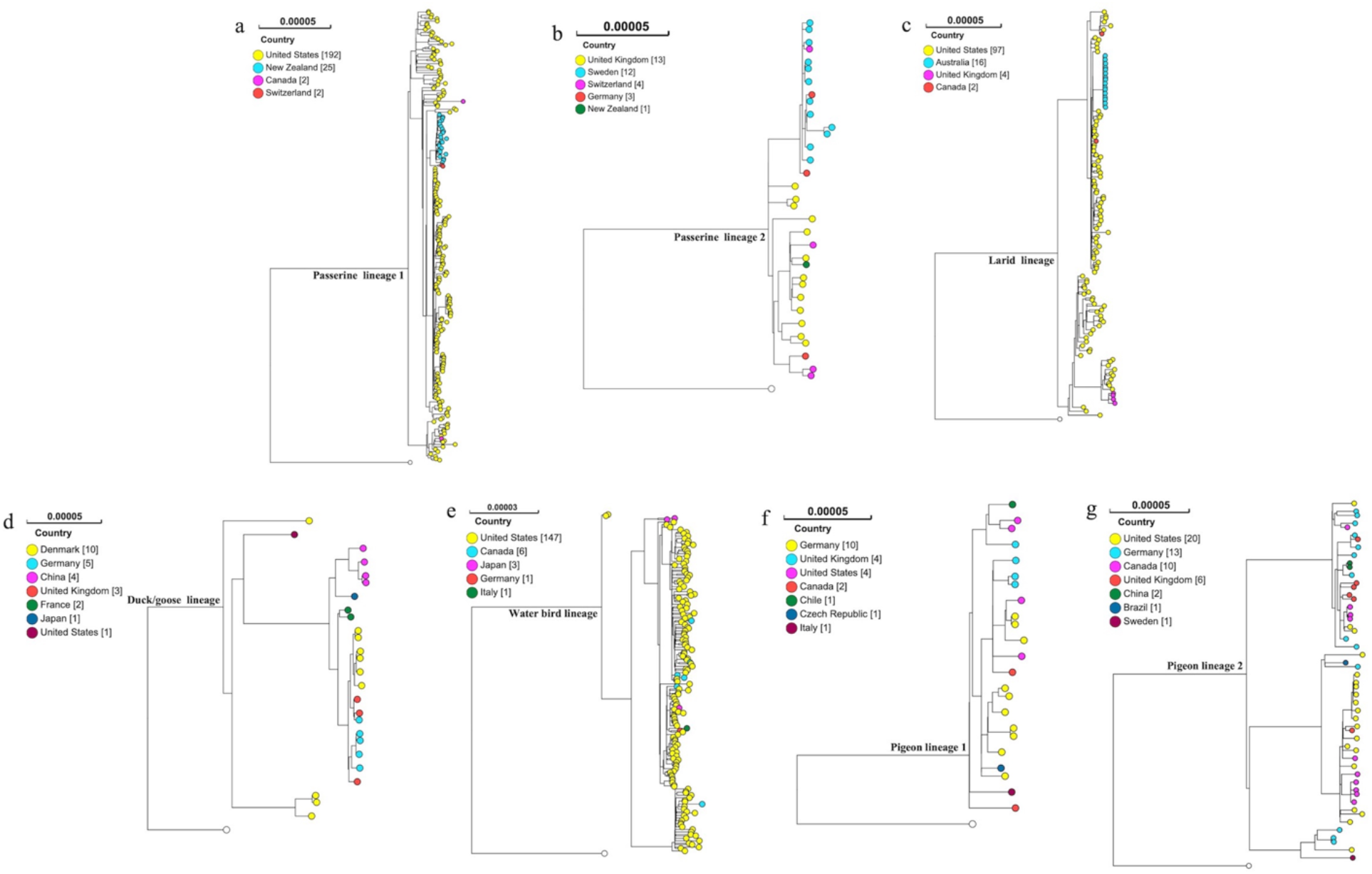
Individual avian host-associated *Salmonella* Typhimurium lineages formed by isolates from different countries. **a,** Passerine lineage 1. **b,** Passerine lineage 2. **c,** Larid lineage. **d,** Duck/goose lineage. **e,** Water bird lineage. **f,** Pigeon lineage 1. **g,** Pigeon lineage 2. Reference genome from *S.* Typhimurium LT2 is represented by white dot in each tree.

**Supplementary Fig. 3:**
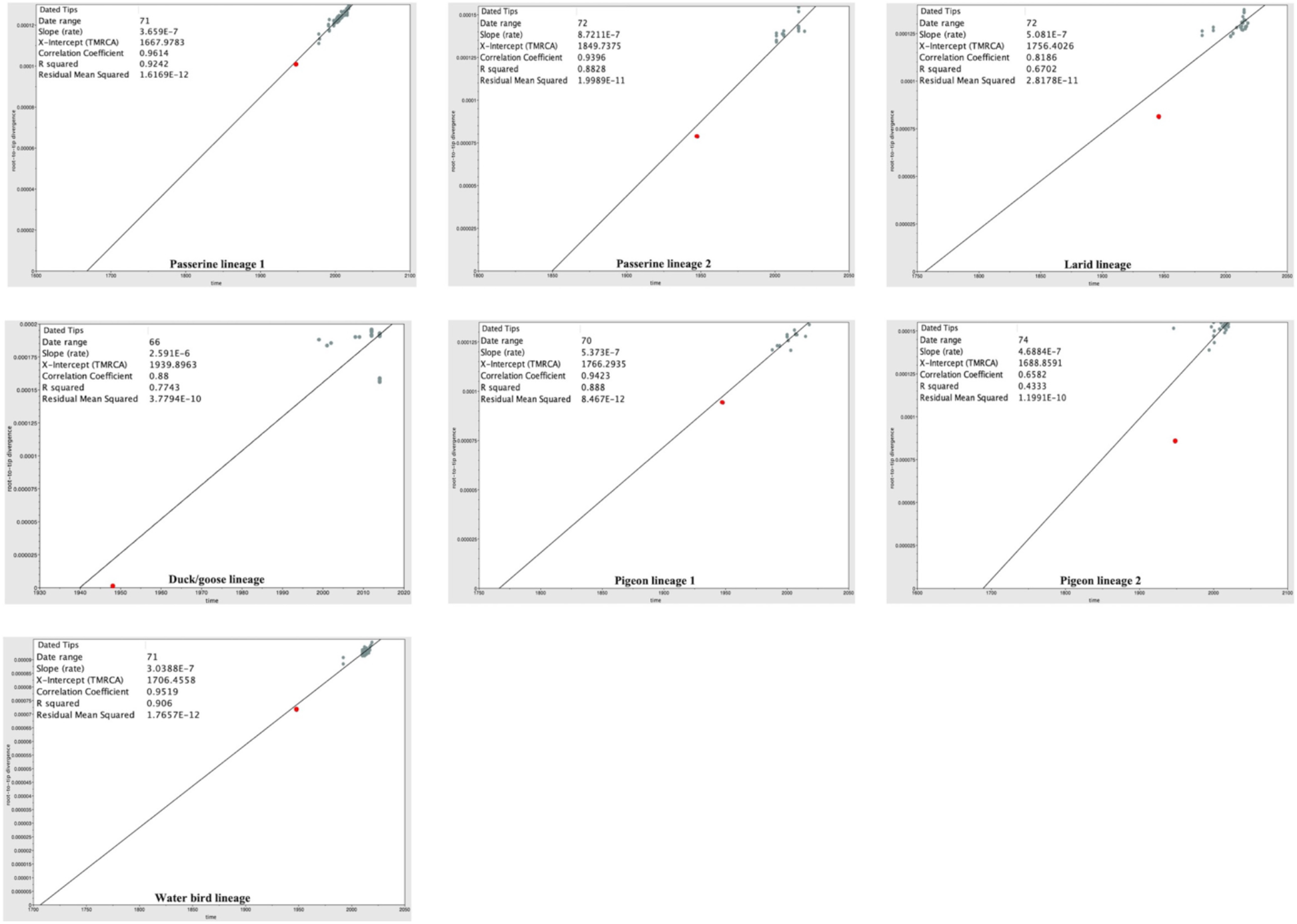
Root-to-tip regression plots showing the temporal signal of the genome sequences used for Bayesian inference. **a,** Passerine lineage 1. **b,** Passerine lineage 2. **c,** Larid lineage. **d,** Duck/goose lineage. **e,** Pigeon lineage 1. **f,** Pigeon lineage 2. **g,** Water bird lineage. Reference genome from *S.* Typhimurium LT2 is represented by red dot.

**Supplementary Fig. 4:**
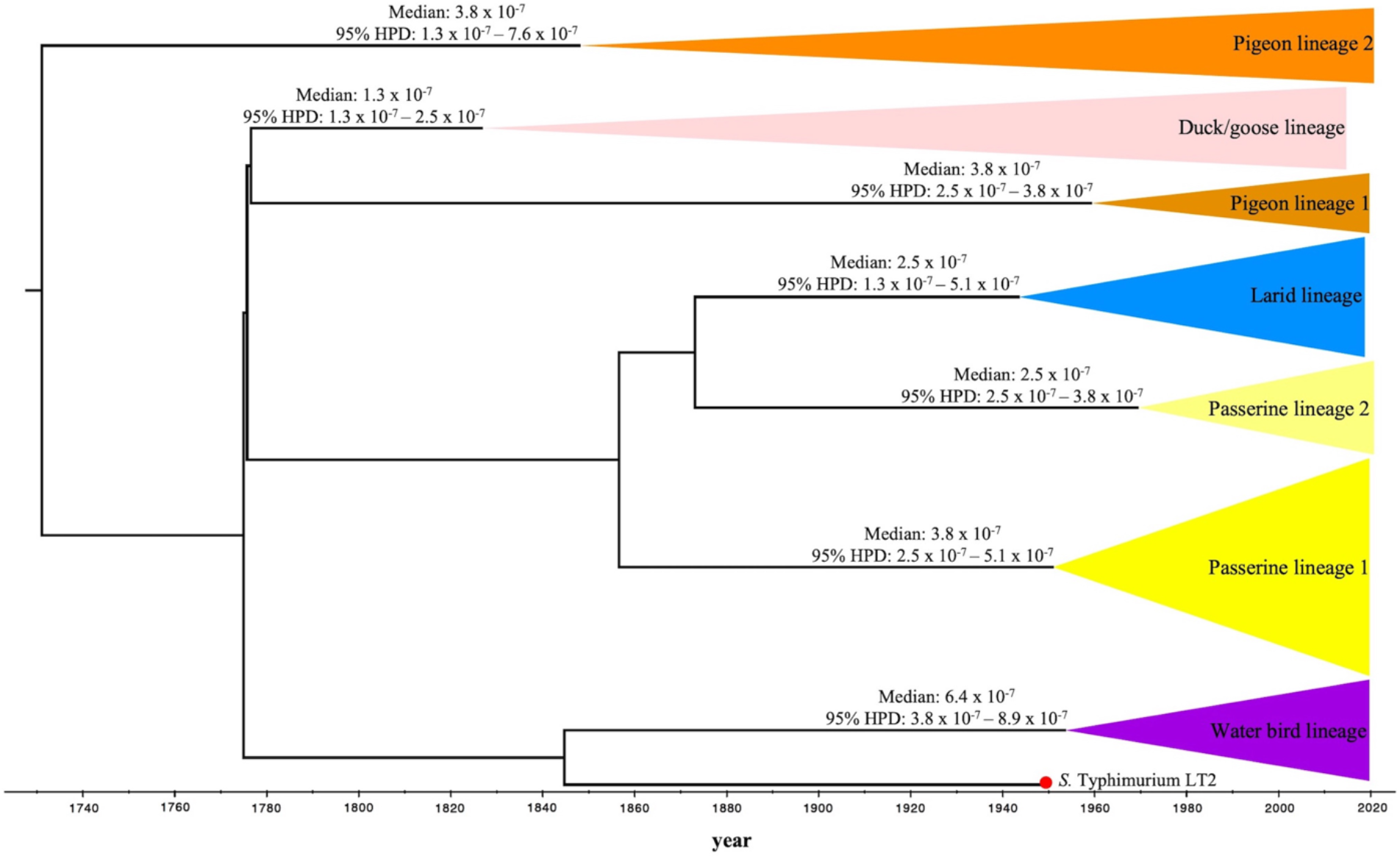
Substitution rates of avian host-associated *Salmonella* Typhimurium lineages inferred by Bayesian time-scaled tree. Estimated substitution rates of individual lineages are reported as median substitution rate with 95% highest posterior probability density (HPD). The red dot at the tree tip represents the reference genome from *S.* Typhimurium LT2 (collection year: ca. 1948). The posterior probability values of representative divergent events are >95% (not shown in the figure).

**Supplementary Fig. 5:**
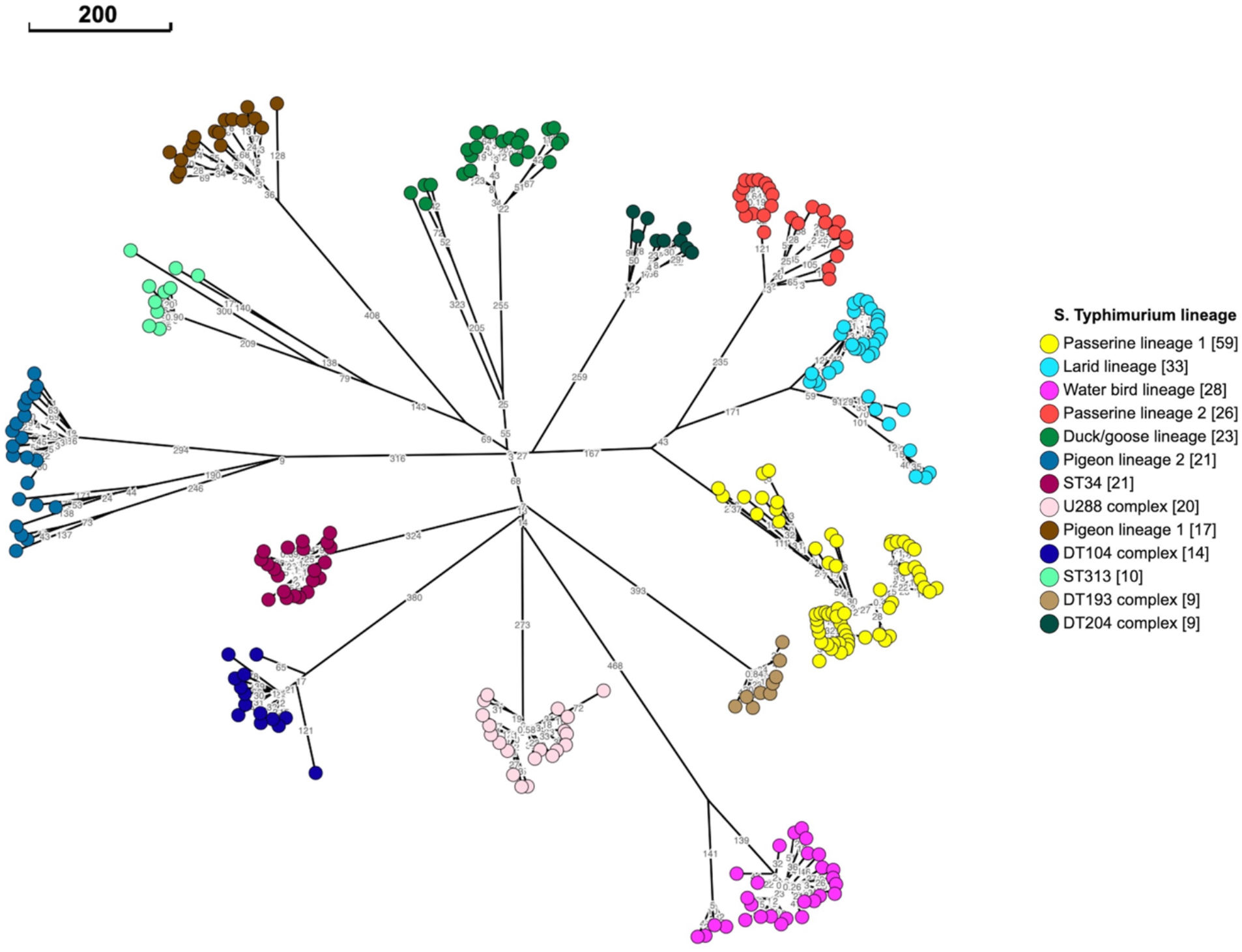
Neighbor joining tree based on wgMLST showing major *Salmonella* Typhimurium lineages circulating in avian (*n* = 207) and non-avian (*n* = 83) host species. Tree tips are colored by *S.* Typhimurium lineage (see key), with number of isolates listed in brackets in the key. The scale bar indicates 200 wgMLST alleles. Allele differences between isolates are indicated by numbers on the connecting lines.

**Supplementary Fig. 6:**
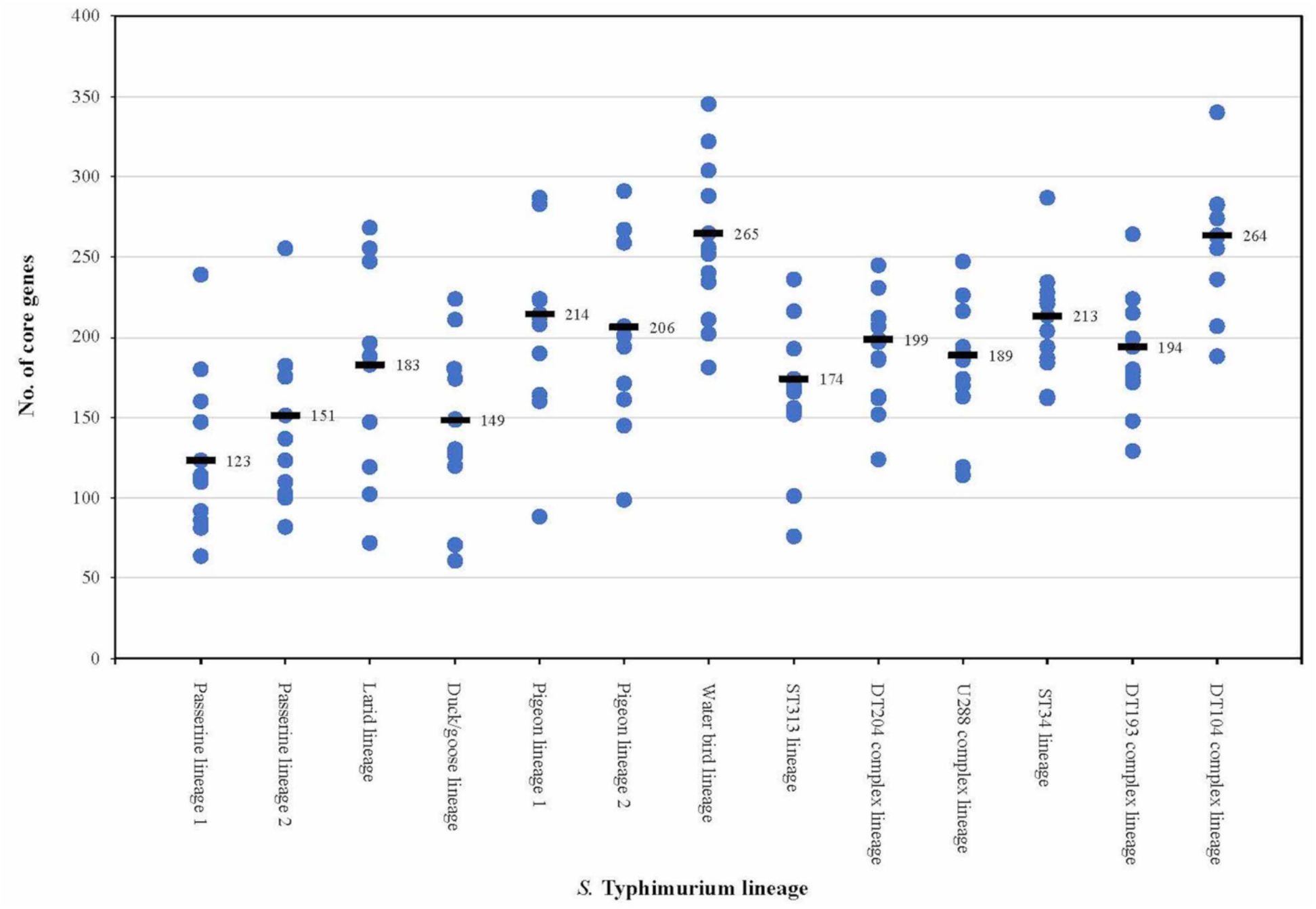
Number of core genes that one *Salmonella* Typhimurium lineage differs another. The blue dot represents the number of core genes that is unique to a specific lineage when comparing it with another lineage. The black line represents the average number of core genes that is unique to a specific lineage when comparing it pairwise with all other lineages. The detailed data of pairwise comparison can be found in Supplementary Data 4.

## Notes

### Competing Interest Statement

The authors have declared no competing interest.

## References

1. Knodler, L. A. & Elfenbein, J. R. Salmonella enterica. Trends Microbiol. 27, 964–965 (2019). https://doi.org/10.1016/j.tim.2019.05.002

2. Stanaway, J. D. et al. The global burden of non-typhoidal salmonella invasive disease: a systematic analysis for the Global Burden of Disease Study 2017. Lancet Infect. Dis. 19, 1312–1324 (2019). https://doi.org/10.1016/S1473-3099(19)30418-9

3. Grimont, P. A. & Weill, F. X. Antigenic formulae of the *Salmonella* serovars. WHO collaborating centre for reference and research on Salmonella 9, 1–166 (2007).

4. Kingsley, R. A. & Bäumler, A. J. Host adaptation and the emergence of infectious disease: the *Salmonella* paradigm. Mol. Microbiol. 36, 1006–1014 (2000). https://doi.org/10.1046/j.1365-2958.2000.01907.x

5. Roumagnac, P. et al. Evolutionary history of *Salmonella* typhi. Science 314, 1301–1304 (2006). https://www.science.org/doi/10.1126/science.1134933

6. Dougan, G. & Baker, S. *Salmonella enterica* serovar Typhi and the pathogenesis of typhoid fever. Annu. Rev. Microbiol. 68, 317–336 (2014). https://doi.org/10.1146/annurev-micro-091313-103739

7. Tanner, J. R. & Kingsley, R. A. Evolution of *Salmonella* within hosts. Trends Microbiol. 26, 986–998 (2018). https://doi.org/10.1016/j.tim.2018.06.001

8. Kingsley, R. A. et al. Genome and transcriptome adaptation accompanying emergence of the definitive type 2 host-restricted *Salmonella enterica* serovar Typhimurium Pathovar. mBio 4, e00565–13 (2013). https://doi.org/10.1128/mBio.00565-13

9. Rabsch, W. et al. *Salmonella enterica* serotype Typhimurium and its host-adapted variants. Infect. Immun. 70, 2249–2255 (2002). https://doi.org/10.1128/IAI.70.5.2249-2255.2002

10. Mather, A. E. et al. Genomic analysis of *Salmonella enterica* serovar Typhimurium from wild passerines in England and Wales. Appl. Environ. Microbiol. 82, 6728–6735 (2016). https://doi.org/10.1128/AEM.01660-16

11. Fu, Y. et al. *Salmonella enterica* serovar Typhimurium from wild birds in the United States represent distinct lineages defined by bird type. Appl. Environ. Microbiol. 88, e01979–21 (2022). https://doi.org/10.1128/AEM.01979-21

12. Didelot, X., Walker, A. S., Peto, T. E., Crook, D. W. & Wilson, D. J. Within-host evolution of bacterial pathogens. Nat. Rev. Microbiol. 14, 150–162 (2016). https://doi.org/10.1038/nrmicro.2015.13

13. Gatt, Y. E. & Margalit, H. Common adaptive strategies underlie within-host evolution of bacterial pathogens. Mol. Biol. Evol. 38, 1101–1121 (2021). https://doi.org/10.1093/molbev/msaa278

14. Bawn, M. et al. Evolution of *Salmonella enterica* serotype Typhimurium driven by anthropogenic selection and niche adaptation. PLoS Genet. 16, e1008850 (2020). https://doi.org/10.1371/journal.pgen.1008850

15. Sheppard, S. K. et al. Niche segregation and genetic structure of *Campylobacter jejuni* populations from wild and agricultural host species. Mol. Ecol. 20, 3484–3490 (2011). https://doi.org/10.1111/j.1365-294X.2011.05179.x

16. Schwartz, D. C. & Cantor, C. R. Separation of yeast chromosome-sized DNAs by pulsed field gradient gel electrophoresis. Cell 37, 67–75 (1984). https://doi.org/10.1016/0092-8674(84)90301-5

17. Hunter, S. B. et al. Establishment of a universal size standard strain for use with the PulseNet standardized pulsed-field gel electrophoresis protocols: converting the national databases to the new size standard. J. Clin. Microbiol. 43, 1045–1050 (2005). https://doi.org/10.1128/JCM.43.3.1045-1050.2005

18. Maiden, M. C. et al. Multilocus sequence typing: a portable approach to the identification of clones within populations of pathogenic microorganisms. Proc. Natl. Acad. Sci. U.S.A. 95, 3140–3145 (1998). https://doi.org/10.1073/pnas.95.6.3140

19. Achtman, M. et al. Multilocus sequence typing as a replacement for serotyping in *Salmonella enterica*. PLoS Pathog. 8, e1002776 (2012). https://doi.org/10.1371/journal.ppat.1002776

20. Callow, B. R. A new phage-typing scheme for *Salmonella* typhimurium. Epidemiol. Infect. 57, 346–359 (1959). https://doi.org/10.1017/S0022172400020209

21. Neoh, H. M., Tan, X. E., Sapri, H. F. & Tan, T. L. Pulsed-field gel electrophoresis (PFGE): A review of the “gold standard” for bacteria typing and current alternatives. Infect. Genet. Evol. 74, 103935 (2019). https://doi.org/10.1016/j.meegid.2019.103935

22. Sheppard, S. K., Guttman, D. S. & Fitzgerald, J. R. Population genomics of bacterial host adaptation. Nat. Rev. Genet. 19, 549–565 (2018). https://doi.org/10.1038/s41576-018-0032-z

23. Schürch, A. C., Arredondo-Alonso, S., Willems, R. J. L. & Goering, R. V. Whole genome sequencing options for bacterial strain typing and epidemiologic analysis based on single nucleotide polymorphism versus gene-by-gene–based approaches. Clin. Microbiol. Infect. 24, 350–354 (2018). https://doi.org/10.1016/j.cmi.2017.12.016

24. Denamur, E. et al. The population genetics of pathogenic *Escherichia coli*. Nat. Rev. Microbiol. 19, 37–54 (2021). https://doi.org/10.1038/s41579-020-0416-x

25. Jarvis, E. D. et al. Whole-genome analyses resolve early branches in the tree of life of modern birds. Science 346, 1320–1331 (2014). https://doi.org/10.1126/science.1253451

26. Prum, R. et al. A comprehensive phylogeny of birds (Aves) using targeted next-generation DNA sequencing. Nature 526, 569–573 (2015). https://doi.org/10.1038/nature15697

27. Fu, Y., Smith, J. C., Shariat, N. W., M’ikanatha, N. M. & Dudley, E. G. Evidence for common ancestry and microevolution of passerine-adapted *Salmonella enterica* serovar Typhimurium in the UK and USA. Microb. Genom. 8, 000775 (2022). https://doi.org/10.1099/mgen.0.000775

28. Fu, Y., M’ikanatha, N. M. & Dudley, E. G. Comparative genomic analysis of *Salmonella enterica* serovar Typhimurium isolates from passerines reveals two lineages circulating in Europe, New Zealand, and the United States.*Appl*. Environ. Microbiol. 88, e00205–22 (2022). https://doi.org/10.1128/aem.00205-22

29. Rambaut, A. Lam, T. T., Max Carvalho, L. & Pybus, O. G. Exploring the temporal structure of heterochronous sequences using TempEst (formerly Path-O-Gen). Virus Evol. 2, 1–7 (2016). https://doi.org/10.1093/ve/vew007

30. Makendi, C. et al. A phylogenetic and phenotypic analysis of *Salmonella enterica* serovar Weltevreden, an emerging agent of diarrheal disease in tropical regions. PLoS Negl. Trop. Dis. 10, e0004446 (2016). https://doi.org/10.1371/journal.pntd.0004446

31. Okoro, C. et al. Intracontinental spread of human invasive *Salmonella* Typhimurium pathovariants in sub-Saharan Africa. Nat. Genet. 44, 1215–1221 (2012). https://doi.org/10.1038/ng.2423

32. Petrovska, L. et al. Microevolution of monophasic *Salmonella* Typhimurium during epidemic, United Kingdom, 2005–2010. Emerg. Infect. Dis. 22, 617–624 (2016). https://doi.org/10.3201/eid2204.150531

33. Mather, A. E. et al. 2013. Distinguishable epidemics of multidrug-resistant *Salmonella* Typhimurium DT104 in different hosts. Science 341, 1514–1517. https://doi.org/10.1126/science.1240578

34. Threlfall, J., Ward, L. R. & Rowe, B. Epidemic spread of a chloramphenicol-resistant strain of *Salmonella* typhimurium phage type 204 in bovine animals in Britain. Vet. Rec. 103, 438–440 (1978). https://doi.org/10.1136/vr.103.20.438

35. Kirkwood, M. et al. Ecological niche adaptation of *Salmonella* Typhimurium U288 is associated with altered pathogenicity and reduced zoonotic potential. Commun. Biol. 4, 1–15 (2021). https://doi.org/10.1038/s42003-021-02013-4

36. Van Puyvelde, S. et al. An African *Salmonella* Typhimurium ST313 sublineage with extensive drug-resistance and signatures of host adaptation. Nat. Commun. 10, 1–12 (2019). https://doi.org/10.1038/s41467-019-11844-z

37. Pulford, CV, et al. Stepwise evolution of *Salmonella* Typhimurium ST313 causing bloodstream infection in Africa. Nat. Microbiol. 6, 327–338 (2021). https://doi.org/10.1038/s41564-020-00836-1

38. Hernandez, S. M. et al. *Epidemiology of a *Salmonella enterica* subsp. enterica* serovar Typhimurium strain associated with a songbird outbreak. Appl. Environ. Microbiol. 78, 7290–7298 (2012) https://doi.org/10.1128/AEM.01408-12

39. Lawson, B. et al. Epidemiological evidence that garden birds are a source of human salmonellosis in England and Wales. PLoS One 9, e88968 (2014) https://doi.org/10.1371/journal.pone.0088968

40. Bloomfield, S. J. et al. Genomic analysis of *Salmonella enterica* serovar Typhimurium DT160 associated with a 14-year outbreak, New Zealand, 1998–2012. Emerg. Infect. Dis. 23, 906–913 (2017). https://doi.org/10.3201/eid2306.161934

41. Ford, L. et al. Whole-genome sequencing of *Salmonella* Mississippi and Typhimurium Definitive Type 160, Australia and New Zealand. Emerg. Infect. Dis. 25, 1690–1697 (2019).https://doi.org/10.3201/eid2509.181811

42. Söderlund, R. et al. Linked seasonal outbreaks of *Salmonella* Typhimurium among passerine birds, domestic cats and humans, Sweden, 2009 to 2016. Euro. Surveill. 24, 1900074 (2019) https://doi.org/10.2807/1560-7917.ES.2019.24.34.1900074

43. Hayden, H. S. et al. Genomic analysis of *Salmonella enterica* serovar Typhimurium characterizes strain diversity for recent U.S. salmonellosis cases and identifies mutations linked to loss of fitness under nitrosative and oxidative stress. mBio 7, e00154 (2016). https://doi.org/10.1128/mBio.00154-16

44. Fu, S. et al. Comparative genomics of Australian and international isolates of *Salmonella* Typhimurium: correlation of core genome evolution with CRISPR and prophage profiles. Sci. Rep. 7, 1–12 (2017). https://doi.org/10.1038/s41598-017-06079-1

45. Salipante, S. J. & Hall, B. G. Determining the limits of the evolutionary potential of an antibiotic resistance gene. Mol. Biol. Evol. 20, 653–659 (2003). https://doi.org/10.1093/molbev/msg074

46. Hiley, L., Graham, R. M. & Jennison, A. V. Genetic characterisation of variants of the virulence plasmid, pSLT, in *Salmonella enterica* serovar Typhimurium provides evidence of a variety of evolutionary directions consistent with vertical rather than horizontal transmission. PloS One 14, e0215207 (2019). https://doi.org/10.1371/journal.pone.0215207

47. McClelland, M. et al. Complete genome sequence of *Salmonella enterica* serovar Typhimurium LT2. Nature 413, 852–856 (2001). https://doi.org/10.1038/35101614

48. Ingle, D. J. et al. Evolutionary dynamics of multidrug resistant *Salmonella enterica* serovar 4,[5],12:i:-in Australia. Nat. Commun. 12, 1–13 (2021). https://doi.org/10.1038/s41467-021-25073-w

49. Bäumler, A. & Fang, F. C. Host specificity of bacterial pathogens. Cold Spring Harb. Perspect. Med. 3, a010041 (2013). https://doi.org/10.1101/cshperspect.a010041

50. Langridge, G. C. et al. Patterns of genome evolution that have accompanied host adaptation in *Salmonella*. Proc. Natl. Acad. Sci. U.S.A. 112, 863–868 (2015). https://doi.org/10.1073/pnas.1416707112

51. Thomson, N. R. et al. Comparative genome analysis of *Salmonella* Enteritidis PT4 and *Salmonella* Gallinarum 287/91 provides insights into evolutionary and host adaptation pathways. Genome Res. 18, 1624–1637 (2008). http://doi.org/10.1101/gr.077404.108

52. Yue, M. et al. Allelic variation contributes to bacterial host specificity. Nat. Commun. 6, 1–11 (2015). https://doi.org/10.1038/ncomms9754

53. Jennings, E., Thurston, T. L. & Holden, D. W. *Salmonella* SPI-2 type III secretion system effectors: molecular mechanisms and physiological consequences. Cell Host Microbe 22, 217–231 (2017). https://doi.org/10.1016/j.chom.2017.07.009

54. Smith, O. M., Snyder, W. E. & Owen, J. P. Are we overestimating risk of enteric pathogen spillover from wild birds to humans? Biol. Rev. 95, 652–679 (2020). https://doi.org/10.1111/brv.12581

55. Muloi, D.M. et al. Population genomics of *Escherichia coli* in livestock-keeping households across a rapidly developing urban landscape. Nat. Microbiol. 7, 581–589 (2022). https://doi.org/10.1038/s41564-022-01079-y

56. Armstrong, G. L. et al. Pathogen genomics in public health. N. Engl. J. Med. 381, 2569–2580 (2019). https://www.nejm.org/doi/10.1056/NEJMsr1813907

57. Perez-Sepulveda, B. M. et al. An accessible, efficient and global approach for the large-scale sequencing of bacterial genomes. Genome Biol. 22, 1–18 (2021). https://doi.org/10.1186/s13059-021-02536-3

58. Fu, Y. et al. Low occurrence of multi-antimicrobial and heavy metal resistance in *Salmonella enterica* from wild birds in the United States. Environ Microbiol. 24, 1380–1394 (2022). https://doi.org/10.1111/1462-2920.15865

59. Timme, R. E. et al. Optimizing open data to support one health: best practices to ensure interoperability of genomic data from bacterial pathogens. One Health Outlook 2, 1–11 (2020). https://doi.org/10.1186/s42522-020-00026-3

60. Zhou, Z. et al. The EnteroBase user’s guide, with case studies on *Salmonella* transmissions, *Yersinia pestis* phylogeny, and *Escherichia* core genomic diversity. Genome Res. 30, 138–152 (2020). https://doi.org/10.1101/gr.251678.119

61. Kumar, S., Stecher, G., Li, M., Knyaz, C. & Tamura, K. MEGA X: molecular evolutionary genetics analysis across computing platforms. Molecular. Biol. Evol. 35, 1547–1549 (2018). https://doi.org/10.1093/molbev/msy096

62. Bouckaert, R. et al. BEAST 2.5: An advanced software platform for Bayesian evolutionary analysis. PLoS Comput. Biol. 15, e1006650 (2019). https://doi.org/10.1371/journal.pcbi.1006650

63. Seemann, T. Shovill: Faster SPAdes assembly of Illumina reads (2017). https://github.com/tseemann/shovill

64. Seemann, T. Prokka: rapid prokaryotic genome annotation. Bioinformatics 30, 2068–2069 (2014). https://doi.org/10.1093/bioinformatics/btu153

65. Page, A. J. et al. Roary: rapid large-scale prokaryote pangenome analysis. Bioinformatics 31, 3691–3693 (2015). https://doi.org/10.1093/bioinformatics/btv421

66. Seemann, T. ABRicate: mass screening of contigs for antibiotic resistance genes (2016). https://github.com/tseemann/abricate

67. Bortolaia, V. et al. ResFinder 4.0 for predictions of phenotypes from genotypes. J. Antimicrob. Chemother. 75, 3491–3500 (2020). https://doi.org/10.1093/jac/dkaa345

68. Carattoli, A. et al. *In silico* detection and typing of plasmids using PlasmidFinder and plasmid multilocus sequence typing. Antimicrob. Agents Chemother. 58, 3895–3903 (2014). https://doi.org/10.1128/AAC.02412-14

69. Liu, B., Zheng, D. D., Jin, Q., Chen, L. H. & Yang, J. VFDB 2019: a comparative pathogenomic platform with an interactive web interface. Nucleic Acids Res. 47, D687–D692 (2019). https://doi.org/10.1093/nar/gky1080

